# Mutations in *rpoB* that confer rifampicin resistance can alter levels of peptidoglycan precursors and affect β-lactam susceptibility

**DOI:** 10.1101/2022.11.14.516545

**Authors:** Yesha Patel, Vijay Soni, Kyu Y. Rhee, John D. Helmann

## Abstract

Bacteria can adapt to stressful conditions through mutations affecting the RNA polymerase core subunits that lead to beneficial changes in transcription. In response to selection with rifampicin (RIF), mutations arise in the RIF resistance determining region (RRDR) of *rpoB* that reduce antibiotic binding. These changes can also alter transcription and thereby have pleiotropic effects on bacterial fitness. Here, we studied the evolution of resistance in *Bacillus subtilis* to the synergistic combination of RIF and the β-lactam cefuroxime (CEF). Two independent evolution experiments led to the recovery of a single *rpoB* allele (S487L) that was able to confer resistance to RIF and CEF through a single mutation. Two other common RRDR mutations made the cells 32x more sensitive to CEF (H482Y) or led to only modest CEF resistance (Q469R). The diverse effects of these three mutations on CEF resistance are correlated with differences in the expression of peptidoglycan (PG) synthesis genes and in the levels of two metabolites crucial in regulating PG synthesis, glucosamine-6-phosphate (GlcN-6-P) and UDP-N-acetylglucosamine (UDP-GlcNAc). We conclude that RRDR mutations can have widely varying effects on pathways important for cell wall biosynthesis, and this may restrict the spectrum of mutations that arise during combination therapy.

**Importance:** Rifampicin (RIF) is one of the most valued drugs in the treatment of tuberculosis. TB treatment relies on a combination therapy, and for multidrug resistant strains may include β-lactams. Mutations in *rpoB* present a common route for emergence of resistance to RIF. In this study, using *B. subtilis* as a model, we evaluate the emergence of resistance for the synergistic combination of RIF and the β-lactam cefuroxime (CEF). One clinically-relevant *rpoB* mutation conferred resistance to both RIF and CEF, whereas two others increased CEF sensitivity. We were able to link these phenotypes to accumulation of specific PG precursors. Mainly, UDP-GlcNAc through its GlmR mediated influence on GlmS activity has a strong impact on CEF resistance. Since these mutations are clinically relevant, these effects on CEF sensitivity may help refine the use of β-lactams in TB therapy.

## Introduction

Bacteria adapt to environmental stresses by coordinated changes in transcription described as bacterial stress responses (1). However, when these phenotypic processes are overwhelmed and the majority of cells are either killed or growth inhibited, there is strong selective pressure for the emergence of adaptive mutations that confer resistance (2). Mutations in *rpoB/rpoC*, encoding the β and β’ subunits of the RNA polymerase (RNAP) core enzyme, can facilitate adaptation to a variety of environmental and antibiotic stresses (3-6). However, the pleiotropic nature of mutations affecting the core RNAP subunits has made it challenging to discern the specific basis of such phenotypes (7). One exception is rifampicin (RIF) resistance (8). RIF binds to the β-subunit of RNAP to suppress transcription, and substitutions in RpoB inhibit RIF-binding, resulting in drug resistance (9). Importantly, such mutations are localized to the RIF-binding pocket and define a RIF-resistance determining region (RRDR). These RRDR changes dramatically reduce RIF-binding, and can also have other less well understood effects on RNA polymerase function (10).

RRDR mutations often have collateral effects, such as reduced growth fitness (11) and altered susceptibility to other antibiotics (12). Accordingly, the ability of RIF to select RNAP mutations has been used as a tool for altering cell physiology (13). One possibility is that the mutant RNAP is altered in its biochemical properties or interactions with regulatory factors, and this leads to a change in the transcriptional landscape. For instance, selection of RIF resistance in *Bacillus subtilis* led to strains defective in sporulation, providing early support for the idea that the genetic program of sporulation might require modifications of RNAP (14). Further, altered expression of metabolic enzymes might account for the effects of *rpoB* mutations on the ability to grow on diverse carbon sources (15). In *Mycobacterium tuberculosis* (MTB), RIF resistant *rpoB* mutants display an altered cell wall metabolism, perhaps due to effect on the channeling of metabolites into cell wall precursors (16, 17).

Since *rpoB* mutations may have global effects on cell physiology, the RRDR mutations that emerge in response to RIF selection can be influenced by other features of the growth environment. This phenomenon has been explored in *B. subtilis*, where both the frequency and spectrum of RRDR mutations is altered in diverse environments (including that of a spaceflight!) (18-20). In a clinical context, RIF is administered as part of a multidrug therapy for the treatment of MTB. Thus, it is important to consider the influence of other antibiotics on acquisition of *rpoB* mutations conferring RIF resistance. More generally, it is important to understand the interactions between co-administered drugs and the impact of evolution of resistance to one drug on susceptibility to the partner drug.

Here, we explore the physiological and genetic interactions between RIF and the cell wall inhibiting β-lactam cefuroxime (CEF). We demonstrate that these two antibiotics are synergistic against *B. subtilis*, and co-selection with these two antibiotics led only to RRDR mutations. However, under these co-selection conditions the spectrum of RRDR mutations was reduced. When commonly arising RRDR mutations were characterized, only one mutation was identified to simultaneously confer high level RIF and CEF resistance. The effects of RRDR mutations on CEF resistance correlated with changes in the expression of peptidoglycan synthesis enzymes and the levels of key intermediates. These findings highlight the ability of RRDR mutations to have divergent effects on microbial physiology.

## Results

### Rifampicin (RIF) and cefuroxime (CEF) exhibit synergy against *B. subtilis*

A synergistic interaction between β-lactams and RIF has been reported against gram-positive bacteria, including both methicillin-resistant staphylococci (21) and mycobacteria (22). We chose to test for synergy in *B. subtilis* between RIF and the cephalosporin cefuroxime (CEF). CEF, with an MIC of 5.12 μg/mL (SI Figure 1A), acts by preferentially binding to and inhibiting the activity of class A PBPs, enzymes involved in the polymerization of PG precursors (23). Using a checkerboard assay, we found the combination of RIF and CEF to be strongly synergistic with a zero interaction potency (ZIP) score (24) of >10 over a range of antibiotic concentrations (Table 1, SI Table 3). Values for the combination of 0.06 μg/mL RIF with increasing concentrations of CEF have been listed in the table for illustration. Full dataset including other concentrations is in SI Table 3. On treatment with sub-MIC concentrations of CEF (up to 0.64 μg/mL), the lag phase was increased by no more than 3 hrs (Figure 1A). A sub-MIC concentration of RIF (0.06 μg/mL) also led to an increase in lag phase (from <1.5 hrs to ∼5 hrs). However, these cells were now very sensitive to growth inhibition by CEF, with as little as 0.08 μg/mL CEF leading to a lag phase of ∼10 hrs (Figure 1B). Similarly, the presence of sub-MIC CEF (0.64 μg/mL) reduced the RIF MIC by 4-fold from 0.125 to 0.03 μg/mL (SI Figure 1B, 1C). This change corresponds to a fractional inhibitory concentration index (FICI) (25) of 0.36, further supporting the conclusion that these two antibiotics act synergistically.

**Table 1:**
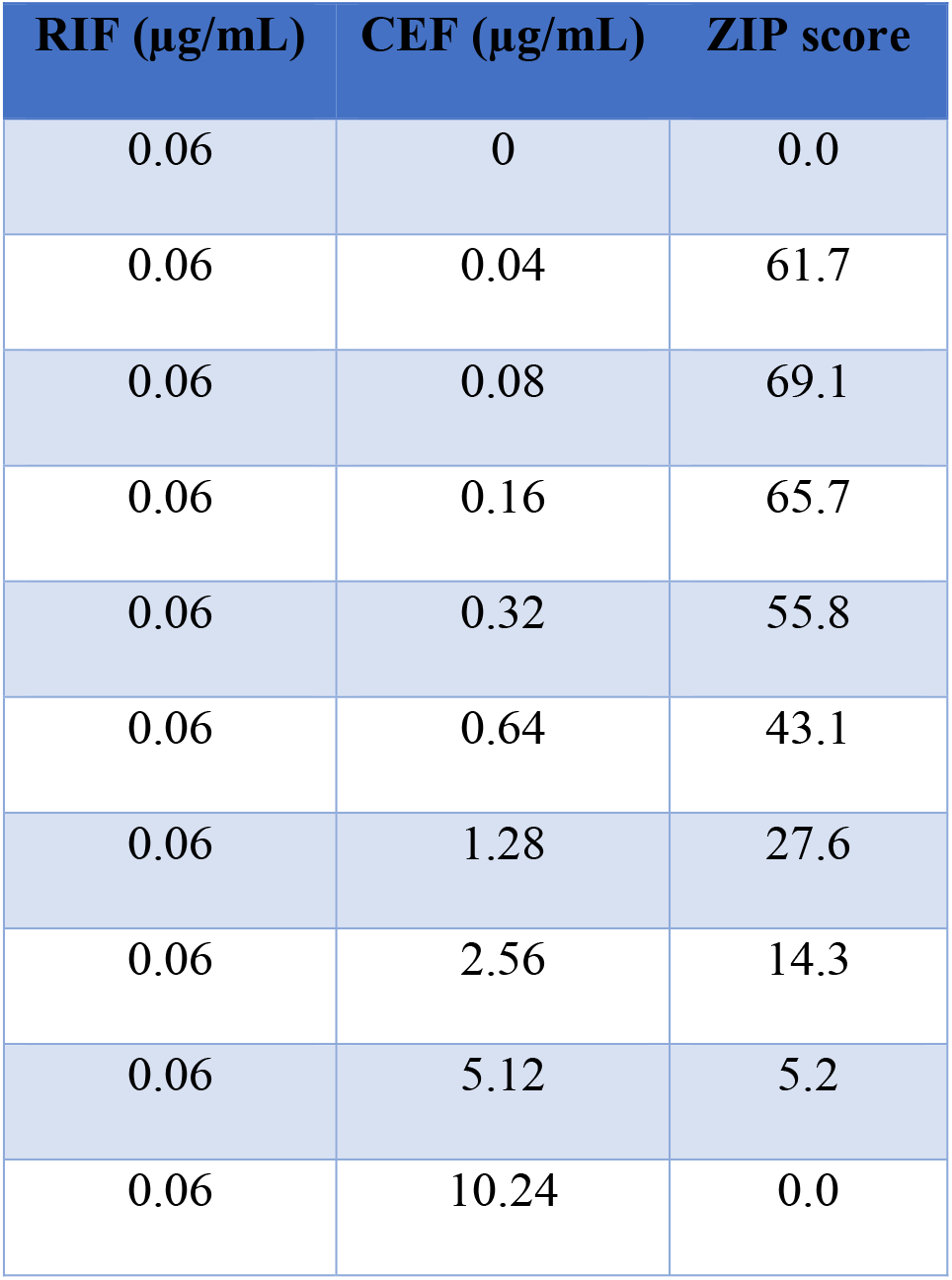
ZIP scores for the combination of 0.06 μg/mL RIF with increasing concentrations of CEF.

| RIF (µg/mL) | CEF (µg/mL) | ZIP score |
| --- | --- | --- |
| 0.06 | 0 | 0.0 |
| 0.06 | 0.04 | 61.7 |
| 0.06 | 0.08 | 69.1 |
| 0.06 | 0.16 | 65.7 |
| 0.06 | 0.32 | 55.8 |
| 0.06 | 0.64 | 43.1 |
| 0.06 | 1.28 | 27.6 |
| 0.06 | 2.56 | 14.3 |
| 0.06 | 5.12 | 5.2 |
| 0.06 | 10.24 | 0.0 |

**Figure 1:**
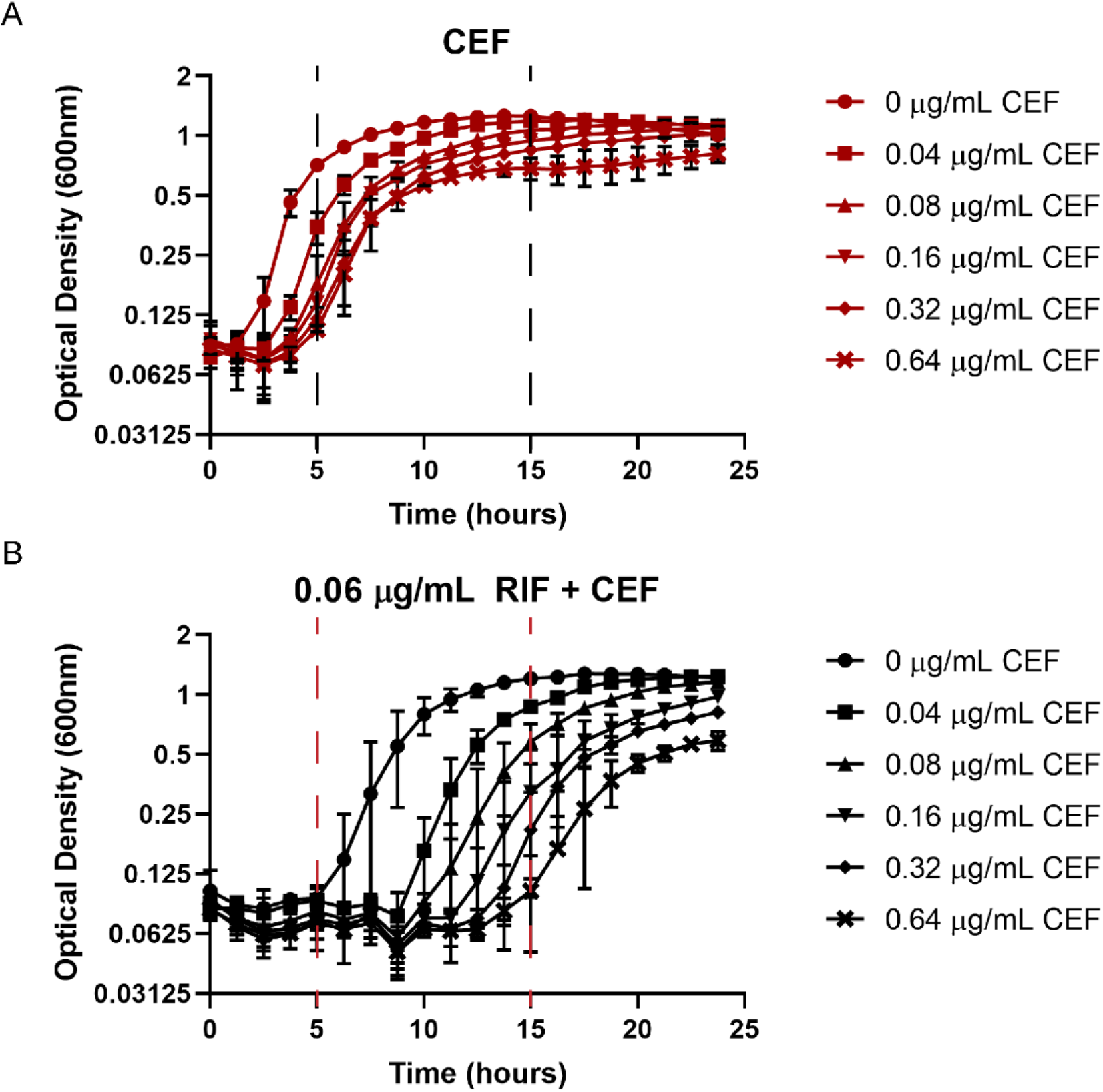
Synergy between rifampicin (RIF) and cefuroxime (CEF) monitored by growth kinetics. Cell density was monitored after treatment with (A) sub-MIC levels of CEF alone, or (B) in the presence of 0.06 μg/mL RIF. The observed lag phases were all less than 5 hr with CEF alone and increased to nearly 15 hr with the combination treatment, as highlighted by the the dashed lines.

### Co-treatment with RIF and CEF selects for mutations in *rpoB*

Drug synergy is a clinically attractive feature of antibiotic chemotherapy. However, drug interactions also have the potential to influence the evolution of resistance (26). Both RIF and CEF susceptibility is influenced by mutations in RNA polymerase (27, 28). We therefore sought to explore how co-treatment with both RIF and CEF affected the evolution of resistance. We hypothesized that the combination of RIF+CEF might select for the emergence of mutations at novel loci. We evolved *B. subtilis* by repeated passage (10x) in the presence of three alternative drug combinations (Figure 2A): 0.06 μg/mL RIF with 2.56 μg/mL CEF (0.5X MIC of the individual drugs), 0.12 μg/mL RIF with 2.56 μg/mL CEF (MIC of RIF and 0.5X MIC of CEF), 0.06 μg/mL RIF with 5.12 μg/mL CEF (0.5X MIC of RIF and MIC of CEF). Under all three conditions, cells developed resistance to both drugs by the fourth passage, as measured by a decrease in diameter in a zone of inhibition (ZOI) assay (Figure 2B-C). The absence of red and blue bars in Figure 2B represents complete loss of ZOI, and hence high resistance to RIF. Interestingly, when evolved in the presence of the highest CEF concentration (5.12 μg/mL), cells were only able to acquire low-level RIF resistance.

**Figure 2:**
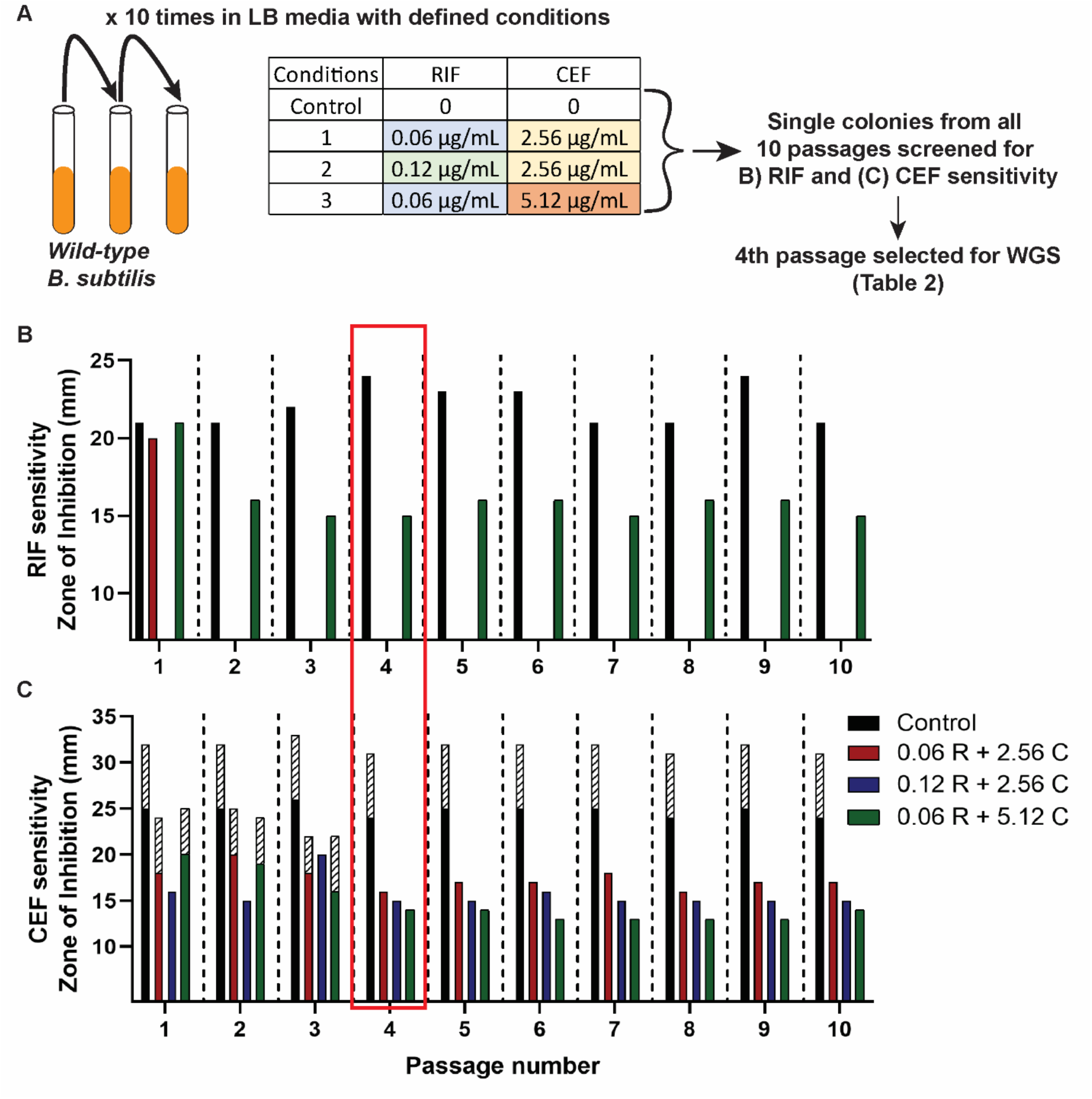
Evolution of WT *B. subtilis* to achieve RIF and CEF resistance. (A) Schematic of the evolution experiment carried out in the presence of the RIF+CEF combination (B) RIF and (C) CEF susceptibilities as measured by zone of inhibition for the 10 passages evolved under the 3 combination treatments. The three RIF and CEF concentrations used for evolving the cells has been mentioned in the legend. The control group consists of 10 passages of WT cells that have not been treated with any drugs. The shaded bars in (C) represent zone of lower density.

To identify the genetic changes associated with resistance, we performed whole genome sequencing (WGS) of single colonies recovered from the 4^th^ passage of selection. Interestingly, all three evolved strains had mutations in the RRDR of *rpoB* (Table 2). This suggests that even in the presence of two drugs, the most facile path to resistance to both drugs is through alterations in the RDDR region of *rpoB*. The two independently evolved strains (A and B) that were selected with sub-MIC levels of CEF both acquired high level RIF resistance with an identical mutation, S487L. The RRDR region is highly conserved (29), and this mutation corresponds to S531L in *E. coli* and S450L, which is the most commonly occurring RIF resistance mutation in *M. tuberculosis* (30). Strain C, evolved with CEF at its MIC (5.12 μg/mL), acquired an *rpoB* P520L mutation that contributed comparatively low level RIF resistance (31). This suggests that the selective pressure imposed by higher CEF concentrations might preclude the acquisition of high RIF resistance through typical RRDR mutations. We sought to confirm this finding by repeating the experiment with ten additional biological replicates. Five tubes were grown with 0.06 μg/mL RIF and 5.12 μg/mL CEF (1x MIC), and five tubes with 0.06 μg/mL RIF and 10.24 μg/mL CEF (2x MIC). In support of the previous experiment, none of the strains acquired high RIF resistance even after 10 passages. Sequencing of the RRDR region from eight isolates led to four strains with atypical RRDR region mutations that led to modest increases in RIF and high CEF resistance (L489S, A478V (2 isolates), S468P), and four that did not contain RRDR mutations. Thus, high levels of CEF seem to impede the emergence of most RRDR region mutations that are known to confer high level RIF resistance in favor of mutations that confer CEF resistance and only partial RIF resistance.

**Table 2:** The mutations identified by WGS after evolution.

| Drug combination | Gene | Coding region change | Amino acid change |
| --- | --- | --- | --- |
| Strain A (0.06R+ 2.56C) | *rpoB* | 1460 C>T | Ser487Leu |
| Strain B (0.12R+ 2.56C) | *rpoB* | 1460 C>T | Ser487Leu |
| Strain C (0.06R+ 5.12C) | *rpoB* | 1559 C>T | Pro520Leu |

### *rpoB* mutants exhibit altered susceptibility to other cell wall acting antibiotics

In addition to characterizing RIF resistant mutants selected by both RIF and CEF (Table 2), we also isolated *rpoB* mutants on agar containing high concentrations (512 μg/mL) of RIF alone. Two additional mutations (H482Y and Q469R) were recovered, which have been identified in prior studies of RIF resistance in *B. subtilis* (32). Mutations in the RRDR residues corresponding to *B. subtilis* S487, H482 and Q469 (Table 3) correspond to more than 90% of RIF resistant MTB clinical isolates. Because of the clinical prevalence of these mutations, and the cross-resistance of the S487L mutant to CEF, we characterized the CEF sensitivity of the H482Y and Q469R RIF resistant mutants (Table 3 and Figure 3A). In contrast to mutants evolved under combination selection (S487L), the H482Y mutation made cells highly susceptible to CEF (32X more sensitive than WT), whereas the Q469R mutation led to a modest increase in CEF resistance (2X more resistant than WT). Combination treatment using RIF and β-lactams has been proposed as a potential drug therapy for *M. tuberculosis* (33). We therefore tested whether two common RIF resistant mutations in *M. tuberculosis* (S531L and H526Y) also alter CEF susceptibility. Indeed, both S531L and H526Y were 2 to 4-fold more sensitive to CEF compared to H37Rv.

**Table 3:** RIF and CEF MICs of different *rpoB* mutants: graded in red for resistance and green for sensitivity compared to WT.

| Mutation<br>(*B. subtilis*) | *E. coli*<br>locus | *M. tuberculosis*<br>locus | RIF MIC<br>(µg/mL) | CEF MIC<br>(µg/mL) |
| --- | --- | --- | --- | --- |
| WT |  |  | 0.125 | 5.12 |
| H482Y | H526Y | H445Y | >4 | 0.16 |
| Q469R | Q513R | Q432R | >4 | 10.24 |
| S487L | S531L | S450L | >4 | 20.48 |
| P520L | P562L | P481L | 4 | 20.48 |

**Figure 3:**
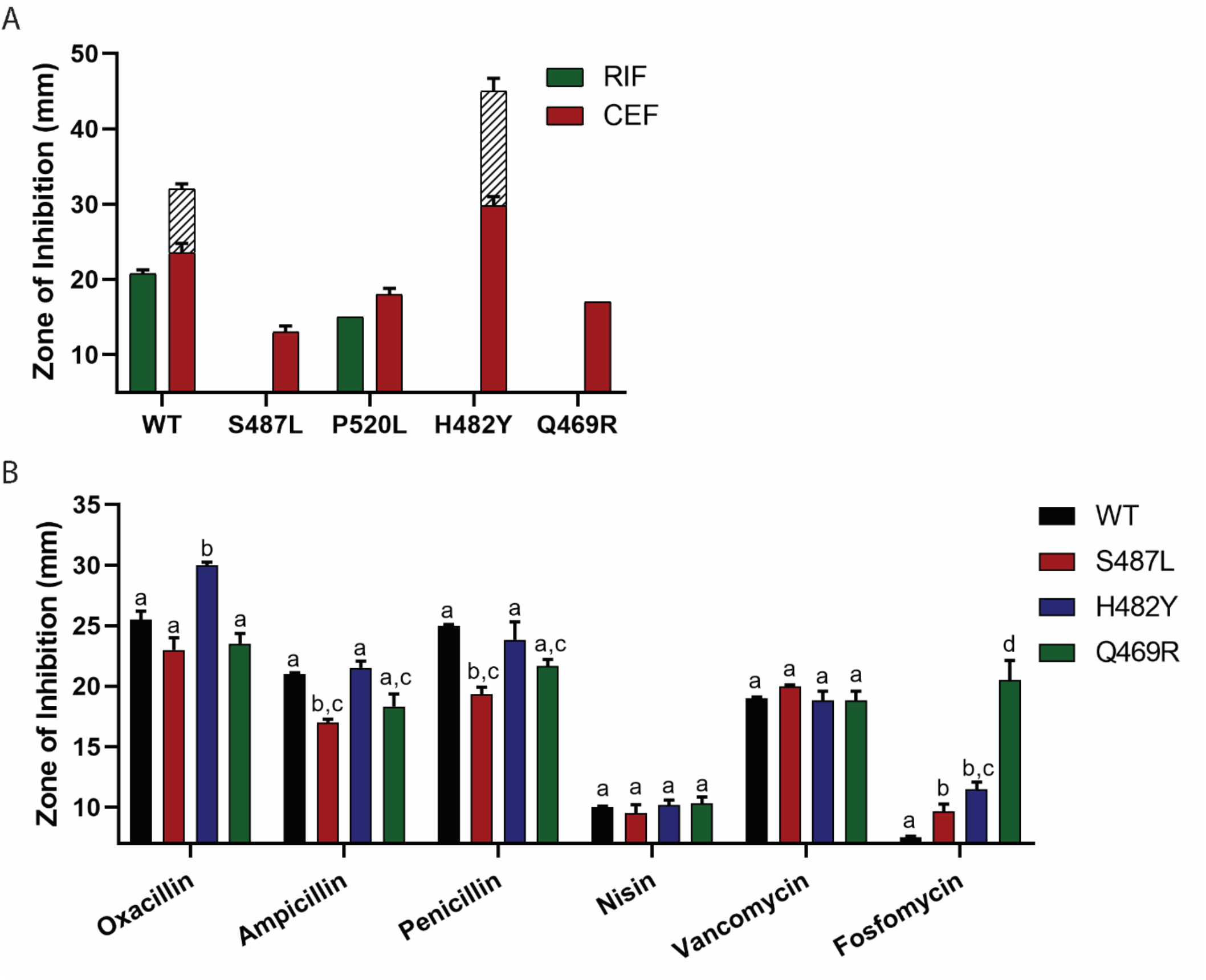
Drug susceptibilities of *rpoB* mutants. (A) Zone of inhibition against RIF and CEF for different *rpoB* mutants (note that only P520L had a detectable inhibition zone with RIF). (B) Zone of inhibition for β-lactams oxacillin, ampicillin and penicillin and other cell wall inhibiting drugs like nisin, vancomycin and fosfomycin for the common clinically associated RIF resistant *rpoB* mutants. The letters indicate the significance of the sensitivity of each strain compared to all others treated with the same antibiotic with *p*-value <0.001 (no comparisons were done between drugs).

Although H482Y frequently emerges in cells subject to RIF selection, this mutation is disfavored in the presence of CEF since it greatly increases CEF sensitivity. Such interactions, where emergence of resistance to one antibiotic increases the susceptibility to another are beneficial in combination therapies (34). We next tested the sensitivity of the three clinically relevant RIF resistance mutants towards additional β-lactams and other antibiotics that target the cell wall (Figure 3B). All β-lactams inhibit the formation of PG layer by targeting different PBPs with varying affinities (35). Three additional β-lactams (oxacillin, ampicillin and penicillin) were similar to CEF with S487L increasing resistance, and H482Y conferring sensitivity. Neither effect was as strong as for CEF, which can be attributed to CEF having the highest affinity for PBP1, the most abundant and primary class A PBP (23).

Extending beyond β-lactams, we also tested the sensitivity of the mutants for nisin and vancomycin, both of which bind lipid II and prevent PG synthesis and crosslinking and, in the case of nisin, can form membrane pores (36, 37). Compared to WT, none of the *rpoB* mutants had a significant difference in sensitivity towards either of these drugs (Figure 3B). In contrast, all the mutants (and especially Q469R) were more susceptible towards fosfomycin (Figure 3B), which inhibits the MurA-dependent synthesis of UDP-N-acetylmuramic acid from UDP-N-acetylglucosamine (UDP-GlcNAc) (38). As a control we also tested the sensitivity of the mutants against drugs acting on other cellular processes: chloramphenicol which inhibits protein synthesis (39); triclosan which inhibits fatty acid synthesis (40) and paraquat which generates ROS toxicity in the cells (41). None of the mutants had a significant difference in the sensitivity against these drugs (SI Figure 2). In conclusion, the predominant *rpoB* mutations associated with high RIF resistance had varying levels of sensitivity to cell wall acting drugs.

### *rpoB* mutations alter the expression of genes affecting PG synthesis

Based on our antibiotic sensitivity results, we hypothesized that these RRDR mutations may change the interaction of RNAP with promoters or regulators involved in the expression of PG synthesis genes. We therefore sought to evaluate the transcript levels of representative PG synthesis genes (*glmS, glmM, glmU, murA* and *ponA*) and two genes that function to divert PG intermediates back into glycolysis (*gamA, nagB*) (Figure 4A). PG synthesis branches from the fructose-6-phosphate node in glycolysis when GlmS converts fructose-6-phosphate to glucosamine-6-phosphate (GlcN-6-P) (42, 43). GlcN-6-P is isomerized by GlmM into GlcN-1-P, which is converted by GlmU to UDP-GlcNAc. MurA initiates synthesis of the second sugar required for PG synthesis, UDP-MurNAc. We included *ponA*, which encodes PBP1, the primary aPBP involved in PG synthesis during vegetative growth and a major target of CEF inhibition (23, 35). PG synthesis can also be supported by import of amino sugars such as GlcNAc present in the growth medium. Catabolism of GlcNAc leads to GlcN-6-P, a branchpoint metabolite that can be used by GlmM to support PG synthesis or when in excess routed into glycolysis through the GamA (44) and NagB (45) enzymes (Figure 4A).

**Figure 4:**
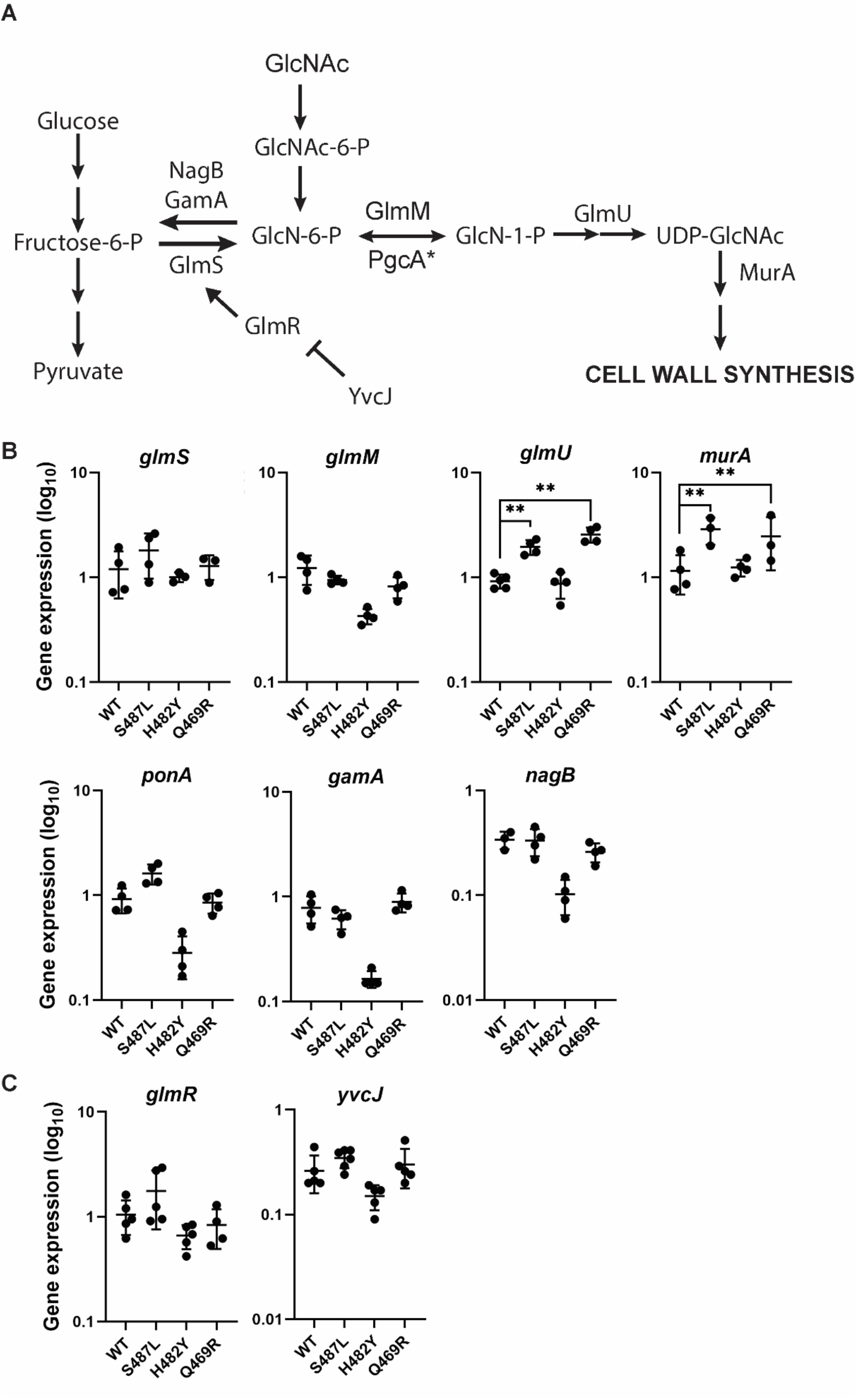
The effect of *rpoB* mutations on peptidoglycan (PG) synthesis. (A) The schematic of PG synthesis pathway. The expression levels of (B) enzymes (C) regulators involved in PG synthesis in WT, and the *rpoB* mutants S487L, H482Y and Q469R as determined by real-time PCR. The expression levels were calculated by the 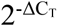 method. *gyrA* was used as the internal control to normalize the levels of the genes of interest. The values are plotted on log10 scale. Significance was calculated using two-way ANOVA with Tukey’s multiple comparisons test. The ** indicates *p*-value less than 0.001.

In the case of the CEF^R^ S487L and Q469R mutants, *glmU* and *murA* were expressed at significantly higher levels compared to WT cells, and other tested genes were unchanged (Figure 4B). CEF resistance was notably not correlated with upregulation of *ponA*, encoding a major target for CEF. In the case of the CEF^S^ H482Y mutant, *glmM* and *ponA* were expressed at lower levels than WT. We hypothesized that reduced *glmM* levels (in the H482Y mutant) might be correlated with an increase in expression of *gamA* and *nagB*. However, mRNA levels of these genes were reduced relative to WT in these mutants. None of the mutants had a difference in their growth kinetics in the absence of any drug (SI Figure 3), suggesting that the altered drug sensitivity of the mutants did not result from slower growth. Thus, we conclude that CEF resistance is correlated with increased transcript levels for some enzymes in PG synthesis (*glmU, murA*), whereas sensitivity is correlated with reduced mRNA levels for other enzymes (*glmM, ponA, gamA, nagB*). Whether these changes in mRNA levels are due to effects of RRDR mutations on RNAP activity at the corresponding promoters or are an indirect effect of other changes in metabolism is not yet clear.

Metabolic flux can be regulated by changes in enzyme activity or enzyme expression. For PG synthesis, GlmS is under complex regulation. The level of *glmS* mRNA is regulated by GlcN-6-P activated mRNA cleavage by the *glmS* ribozyme (46). However, the mRNA level of *glmR* was little changed in the RRDR mutants relative to WT (Figure 4C). In addition, GlmS activity is allosterically activated by GlmR (47). GlmR activity is antagonized by complex formation with YvcJ in the presence of high UDP-GlcNAc (48). Therefore, we sought to determine whether RRDR mutations affect the levels of metabolites that might impact PG synthesis.

### RRDR mutations alter the levels of key PG intermediates

To monitor the impact of RRDR mutations on metabolite pools we performed untargeted metabolomics. We focused our attention of the levels of the two key regulatory intermediates noted above, GlcN-6-P and UDP-GlcNAc (Figure 4A), and pyruvate, which is indicative of the flux of fructose-6-phosphate into glycolysis (49). The Q469R strain did not show any significant difference in the levels of these metabolites, so we focused on the differences between the CEF^R^ (S487L) and CEF^S^ (H482Y) strains (Figure 5).

**Figure 5:**
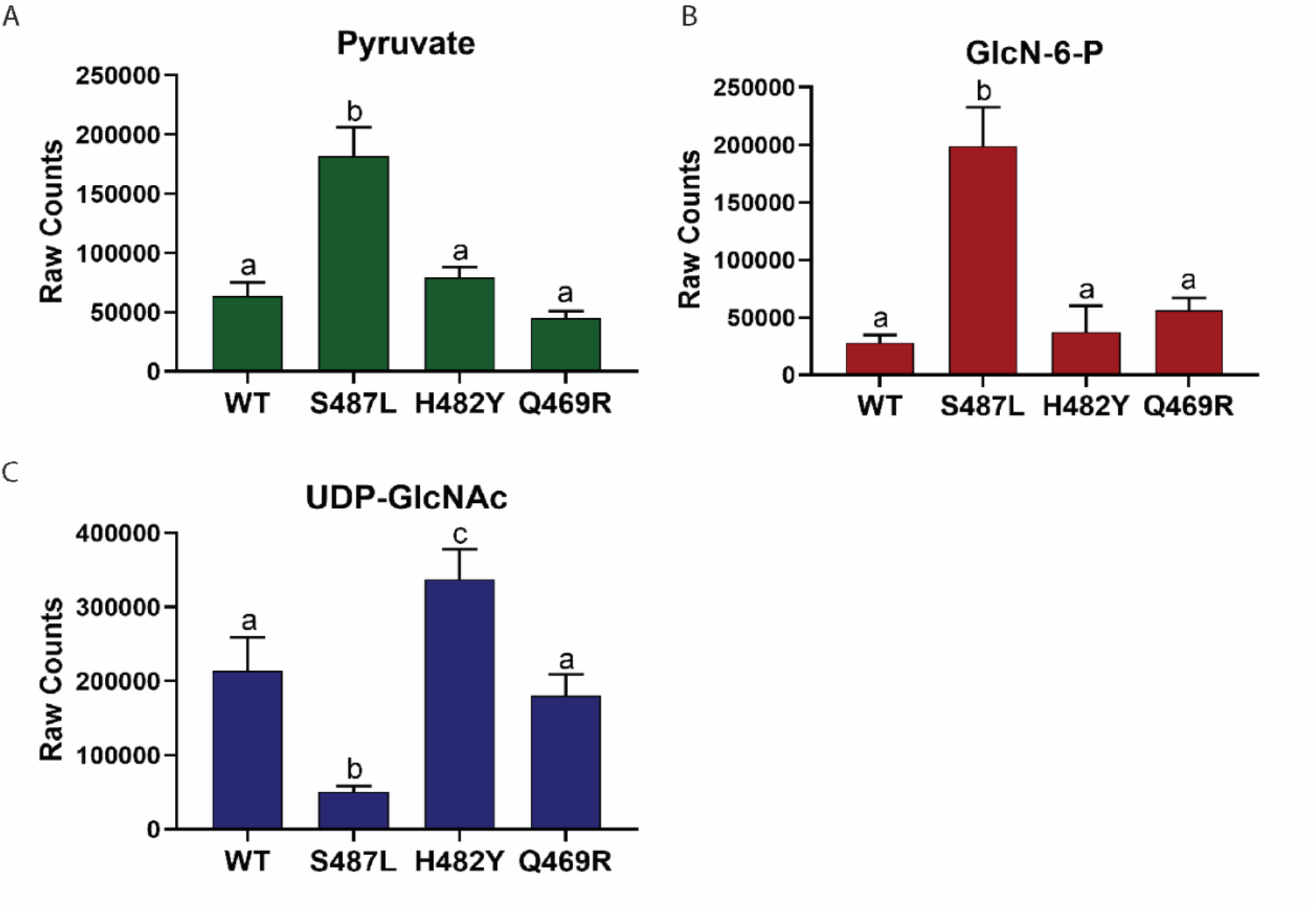
Metabolite levels in *rpoB* mutants. (A) Metabolite levels of pyruvate which indicate the flux through glycolysis (B) Metabolite levels of GlcN-6-P which determine flux into PG synthesis (C) Metabolite levels of UDP-GlcNAc which indicate the rate of PG formation. Same letters define dataset with no significant difference. Different letters define a significant difference of *p*-value less than 0.0001 amongst the mutants.

For the CEF^R^ S487L mutant, we observed an increase in GlcN-6-P and a decrease in UDP-GlcNAc. Since UDP-GlcNAc regulates GlmS activity through the YvcJ/GlmR pathway (Figure 4A), low UDP-GlcNAc will lead to high GlmS activity, which might account for elevated GlcN-6-P. We also noted elevated mRNA levels for *glmU* and *murA* (Figure 4B). Thus, we conclude that the S487L mutant has changes in both gene expression and metabolite levels consistent with a higher rate of PG synthesis. Although one might expect that elevated GlcN-6-P could reduce *glmS* mRNA levels (by ribozyme cleavage) and increase the expression of *gamA* and *nagB*, our qRT-PCR results showed no evidence for these changes (Figure 4B), suggesting that GlcN-6-P has not reached levels needed to trigger these responses.

In contrast, the CEF^S^ H482Y mutant had elevated levels of UDP-GlcNAc. In this case, we predict that the high UDP-GlcNAc will cause sequestration of GlmR in a YvcJ:GlmR:UDP-GlcNAc complex and thereby prevent GlmR stimulation of GlmS activity (48). By restricting GlmS activity, this could reduce flux of fructose-6-P into PG and contribute to the CEF sensitive phenotype. Thus, the most striking correlation to emerge from the metabolomics analysis is the correlation between UDP-GlcNAc and CEF sensitivity. Further, our data support the idea that a key function of UDP-GlcNAc is as a feedback regulator of GlmS activity, as mediated by the GlmR/YvcJ pathway (48).

### The ability of UDP-GlcNAc to modulate PG synthesis is dependent on GlmR

We used epistasis studies to determine if the correlation of UDP-GlcNAc levels and CEF sensitivity is in fact mediated by the role of UDP-GlcNAc as a negative regulator of GlmR activity. The CEF^R^ S487L mutant has reduced UDP-GlcNAc levels that could result in increased activity of the GlmR regulator and this, in turn, could lead to elevated PG synthesis and contribute to antibiotic resistance. Consistent with this model, the elevated CEF^R^ of the S487L mutant is lost in a strain additionally lacking *glmR* (Figure 6). Conversely, in the CEF^S^ H482Y mutant UDP-GlcNAc levels are high, and therefore we predict that GlmR will be largely non-functional due to sequestration in a YvcJ:GlmR:UDP-GlcNAc complex (48). Both the H482Y and the *glmR* mutations individually make cells CEF^S^ but these two mutations are not additive in the H482Y *glmR* double mutant (Figure 6.). This supports our hypothesis that H482Y and *glmR* function in the same pathway, and that H482Y has effectively inactivated GlmR function by altering metabolism leading to a high level of UDP-GlcNAc.

**Figure 6:**
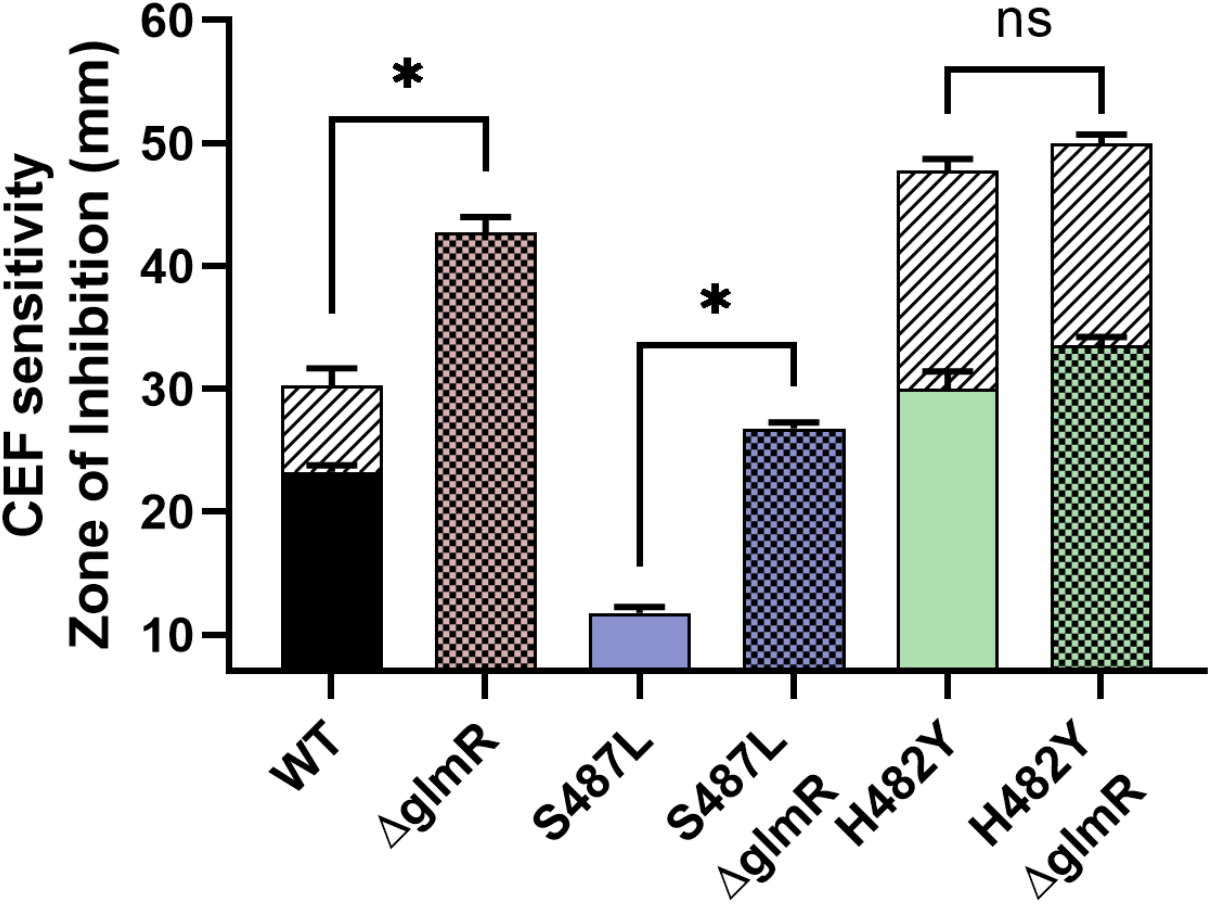
The importance of GlmR activity in CEF sensitivity. The sensitivity of WT and *rpoB* mutants with and without the deletion of *glmR* as measured by zone of inhibition. An * indicates *p*-value less than 0.0001.

### Perturbing flux of amino sugars can alter CEF sensitivity

We hypothesize that the CEF sensitivity of the H482Y mutant is due to restricted GlmS activity resulting from elevated UDP-GlcNAc levels. Therefore, we sought to bypass GlmS by supplementing cells with GlcNAc, which has been shown to increase the level of GlcN-6-P (50). Indeed, in the presence of GlcNAc there was a significant increase in CEF resistance for the H482Y mutant (Figure 7A). The growth of the cells in liquid media in the presence of 0.04 μg/mL of CEF was also significantly better when LB was supplemented with GlcNAc (SI Figure 4). These results suggest that increasing flux of sugars into PG synthesis restores CEF resistance to H482Y by bypassing GlmS. Consistently, if we instead delete *gamA* (Figure 4A) the flux of amino sugars present in the growth medium into glycolysis is restricted, and this also increases CEF resistance. We next tested the impact of increasing the flow of GlcNAc into UDP-GlcNAc on CEF resistance. We ectopically induced expression of the GlmM phosphoglucosamine mutase (PNGM) and PgcA*, an allele of phosphoglucomutase with increased PNGM activity (51). Neither gene was able to increase CEF resistance (Figure 7B). This is consistent with the hypothesis that GlmS activity is restricted, GlcN-6-P is a limiting metabolite for PG synthesis, and only the import of amino sugars from outside the cell can bypass this restriction.

**Figure 7:**
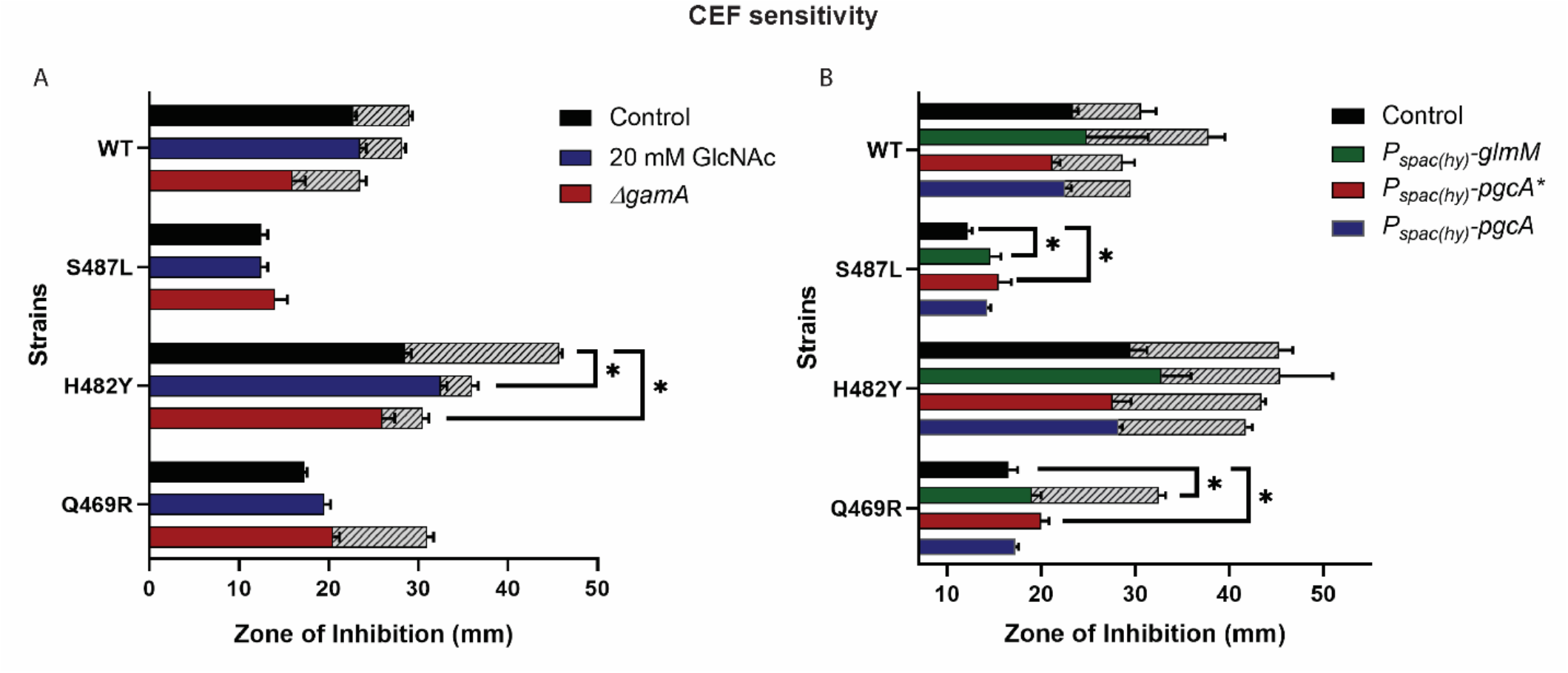
Perturbation of GlcN-6-P and UDP-GlcNAc levels in the cells. The sensitivity of WT and *rpoB* mutants against CEF as measured by zone of inhibition on (A) media supplemented with 20 mM GlcNAc and on deletion of *gamA* which directs GlcN-6-P towards glycolysis (B) on induction of the phosphoglucosaminemutase *glmM* and *pgcA*\* and phosphoglucomutase *pgcA*. An * indicates *p*-value less than 0.01.

Conversely, the CEF^R^ S487L mutant did not exhibit any difference in CEF sensitivity in the presence or absence of 20 mM GlcNAc, or upon deletion of *gamA* (Figure 7A). This is consistent with our hypothesis that this strain is not restricted in the flux of F6P into GlcN-6-P. In this case, induction of *glmM* or *pgcA*\* actually led to a slight increase in CEF sensitivity. In contrast, induction of PgcA, which has comparatively low PNGM activity (51), had no effect. We speculate that with this strain, which has high GlcN-6-P levels (Figure 7B), further increase in synthesis of amino sugars leads to a metabolic imbalance. Finally, for the Q469R mutant, which did not exhibit any significant depletion or accumulation of the PG intermediates, GlcNAc addition did not change CEF susceptibility. Similar to S487L, induction of *glmM* or *pgcA*\* in Q469R also led to a slight increase in CEF sensitivity (Figure 7B).

## Discussion

Drug interactions have a strong impact on evolution of resistance (52). Here, we evaluated the emergence of resistance to a combination of a β-lactam (CEF) and rifampicin (RIF). These two drugs are synergistic in *B. subtilis*, as shown also for other bacteria (53-56). We used *in vitro* evolution followed by whole-genome sequencing to identify mutations that enable growth in the presence of this dual selection. Strikingly, only one single RRDR mutation (S487L) emerged that confers high level resistance to both antibiotics. With CEF at or above the MIC, the acquisition of high-level RIF resistance was restricted. When this selection was repeated and colonies screened specifically for RRDR mutations we identified several other mutations, not commonly associated with RIF resistance, that confer high level CEF resistance and only modestly increase RIF resistance.

These results highlight the importance of RRDR mutations in RIF resistance (by reducing RIF binding to the β-subunit), and the ability of *rpoB* mutations to also confer resistance to other antibiotics by less direct mechanisms. In the presence of CEF, only a limited set of mutations can simultaneously lead to CEF and RIF resistance, and these were found in the RRDR. In fact, other common RRDR mutations that confer high level RIF resistance were either sensitive (H482Y) or had lower resistance to CEF (Q469R). In MTB both mutants corresponding to S487L and H482Y were sensitive to CEF compared to WT. The collateral sensitivity to CEF on acquiring RIF resistance is favorable when considering multidrug treatment (57). Further, co-treatment with β-lactams and RIF may constrain emergence of RIF resistance.

Previously, *rpoB* mutations have been described that alter susceptibility to cell wall inhibiting drugs such as β-lactams (6, 27), vancomycin, and daptomycin (58). However, resistance mutations directly selected by each of these drugs typically do not map to RRDR (6). However, some RIF resistance mutations in the RRDR not only decrease RIF binding, but can lead to alterations in the cell wall (16). In MTB, the frequently occurring H526Y RRDR mutant is very sensitive to cell wall inhibitors, and to the deletion of genes encoding auxiliary functions related to cell wall synthesis and division (59). Similarly, we report here that *B. subtilis* RRDR mutations can lead to either sensitivity or resistance to an antibiotic (CEF) that inhibits PG synthesis.

The identification of S487L (CEF^R^) and H482Y (CEF^S^) mutants in *B. subtilis* presents a useful tool to understand the impact of RRDR mutations on cell wall homeostasis. Using transcriptomic and metabolomic studies, we present evidence for the importance of altered metabolite levels (GlcN-6-P and UDP-GlcNAc) in affecting β-lactams susceptibility. Specifically, higher levels of UDP-GlcNAc in H482Y are correlated with CEF sensitivity, which we ascribe to a loss of GlmR-mediated activation of GlmS. Metabolic feeding studies and genetic epistasis suggests this to be a direct cause of the altered resistance. Conversely, the S487L mutant maintains high levels of GlcN-6-P and low levels of UDP-GlcNAc, and in this strain GlmR-mediated activation of GlmS is critical for maintaining PG synthesis. Although not the intent of this study, our results have served to highlight the importance of GlmR as a key regulator of metabolic flux through GlmS, the enzyme that shunts carbon from glycolysis/gluconeogenesis into amino sugar and PG synthesis. Drugs that inhibit PG synthesis cause a buildup of cell wall intermediates including UDP-GlcNAc (60). When UDP-GlcNAc levels increase, it binds to GlmR and flux into PG synthesis may be reduced. Since GlmR is conserved in many bacteria, including MTB (61, 62), these types of effects are important to consider when considering mechanisms of adaptation and resistance to cell wall antibiotics.

Here, we have validated the central role of GlmR as a regulator, and UDP-GlcNAc as a regulatory metabolite, using the divergent effects of the S487L and H482Y RRDR mutations on CEF resistance. We have used three experimental perturbations to alter the availability of metabolites to support PG synthesis: GlcNAc supplementation and restriction of catabolism (*gamA* deletion), elevated expression of *glmM* or *pgcA*\*, and deletion of *glmR* (Figure 8). The CEF^R^ S487L mutant maintains high levels of GlcN-6-P and low levels of UDP-GlcNAc. Thus, in this strain GlmR is active and maintains relatively higher flux of PG synthesis and is better able to tolerate high levels of CEF. As predicted, if GlmR is deleted the cells become more sensitive to CEF (Figure 6). Due to the high levels of GlcN-6-P in this strain, induction of *glmM* and *pgcA*\*, combined with elevated *glmU* expression (Figure 4), may lead to elevation of UDP-GlcNAc. High UDP-GlcNAc, in turn, will inactivate GlmR, restrict flux to PG synthesis, and increase CEF sensitivity (Figure 7B, 8). In the case of CEF^S^ H482Y, the cells already have high levels of UDP-GlcNAc, which restricts PG synthesis and confers CEF sensitivity. The only manipulation that increased the resistance of this strain was supplementation with GlcNAc or *gamA* deletion, both of which increase PG synthesis independent of GlmR (Figure 7A, 8).

**Figure 8:**
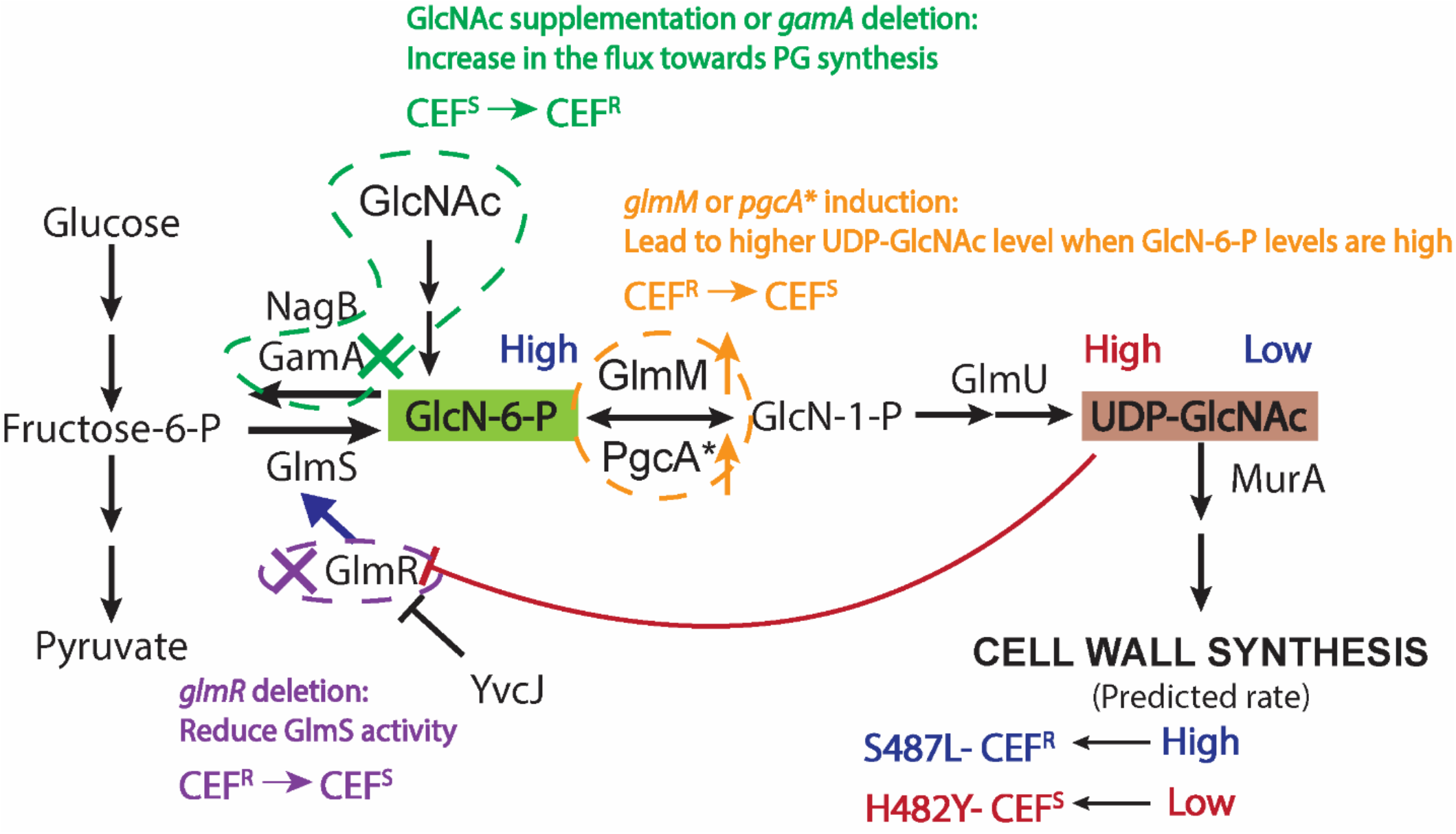
Interpretation of experimental perturbations predicted to affect UDP-GlcNAc levels. The text and arrow in blue summarize the data for the CEF^R^ S487L mutant. This mutant has low levels of UDP-GlcNAc. Thus, GlmR is free to stimulate GlmS activity and cells are predicted to maintain a high rate of PG synthesis detected by high levels of GlcN-6-P. The text and arrow in red summarize the data for CEF^S^ H482Y mutant. This mutant has high levels of UDP-GlcNAc which would bind with GlmR. The bound GlmR is unavailable to stimulate GlmS activity and is thereby predicted to reduce PG synthesis. Three perturbation scenarios are also presented. In green, cells were supplemented with 20 mM GlcNAc or *gamA* was deleted. Both led to higher flux towards PG synthesis independent of GlmS thereby bypassing the bottleneck in the H482Y mutant and leading to elevated CEF resistance. In orange, induction of *glmM* or *pgcA*\* is predicted to increase the levels of UDP-GlcNAc, but only in S487L which has high levels of GlcN-6-P. Thus, this treatment is predicted to block GlmR-dependent GlmS activation in S487L, reduce PG synthesis, and thereby contribute to CEF sensitivity. These inferences are supported by analysis of the effects of a *glmR* deletion (purple). In S487L we observed low UDP-GlcNAc levels and predict that GlmR is activating GlmS. Consistently, deletion of *glmR* makes the S487L strain more CEF sensitive. In contrast, in H482Y we predict that the high observed UDP-GlcNAc levels will keep GlmR sequestered in an inactive state, and consistently there is no effect of deleting *glmR*.

In summary, we have shown that RIF^R^ RRDR mutants have altered susceptibility to β-lactams due to altered levels of PG metabolites, especially UDP-GlcNAc. RRDR mutations have a global impact on the transcriptome of the cells and can lead to pleiotropic effects. Analyzing the expression levels of PG synthesis genes did not reveal why UDP-GlcNAc levels are altered by RRDR mutations, but the downstream effects of this altered metabolite can account for differences in sensitivity to β-lactams. β-lactams are some of the most powerful antibiotics and are being considered in TB therapy with RIF (33). Thus, this work on evolution of resistance to the combination of RIF and CEF, the collateral sensitivity to CEF on acquisition of RIF resistance, and the differential response of *rpoB* mutants to CEF will benefit future studies designing effective drug treatments.

## Materials and Methods

### Bacterial strains, plasmids and growth conditions

Bacterial strains used in this study are listed in SI Table 1. All stains were grown in lysogeny broth (LB) medium at 37°C. Liquid cultures were aerated on an orbital shaker at 280 rpm. Glycerol stocks were streaked on LB agar plates and incubated overnight at 37°C. *rpoB* was amplified using the primers mentioned in SI Table 2. Mutations in the RIF-resistance determining region (RRDR) of *rpoB* were confirmed by Sanger sequencing at the Biotechnology Resources core facility at Cornell University using primer 9286. *glmR::erm* and *gamA*::*erm* were ordered from the BKE collection available at the Bacillus Genetic Stock Centre (BGSC) (63). The gene deletion with the erythromycin cassette was then transformed into the desired strains by natural competence induced in modified competence (MC) medium. The cassette was removed using pDR244 as described previously (63). Transformation was done using chromosomal DNA with selection on plates having 1 μg/mL of erythromycin and 25 μg/mL lincomycin. The deletion was confirmed by PCR with check primers listed in Supplementary Table 2. Strains with inducible expression of *glmM* (HB16910), *pgcA*\* (HB16946) and *pgcA* (HB16945) were made using chromosomal DNA from strains from a previous study (51). Genes were ectopically expressed at *amyE* locus under promoter P_*spac(hy)*_ and selection of transformants was performed in the presence of chloramphenicol (10 μg/mL)

### Growth kinetics and MIC determinations

A bioscreen C growth curve analyzer (Growth curves USA, NJ) was used to monitor the growth of the strains. Initially, cultures were grown up to ∼0.4 OD_600_ in 5 mL culture tubes. 1 μL of this culture was inoculated in each well of honeycomb 100 well plates containing 200 μL of LB media. OD_600_ was monitored every 15 min up to 24 hrs with constant shaking at 37°C. For MIC determination, two-fold increase in the drug concentration was screened ranging from 0.04 to 10.24 μg/mL for CEF and 0.075 to 4 μg/mL for RIF. The minimum concentration of drug having at least 90% growth inhibition compared to the untreated control after 8 hrs of treatment was considered as MIC for the drug. Control cells reach stationary phase within 8 hrs (OD_600_ ∼ 1.0). Percent inhibition was calculated as follows:

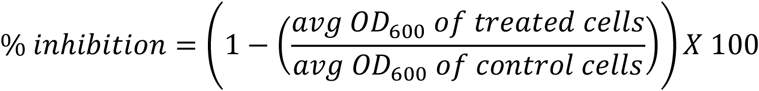

Average OD_600_ was calculated from 3 biological replicates.

### Synergy quantification

Checkerboard assays were used to determine the interaction between RIF and CEF (64) with 2-fold dilutions of both drugs. 1 μL of 0.4 OD cultures was added in each well of 200 μL media containing either or both drugs. The MIC of the drug combination was determined as mentioned in the previous section. To quantify the interaction between the two drugs we calculated both a fractional inhibitory concentration index (FICI) and a ZIP score. The formula to calculate FICI is as follows:

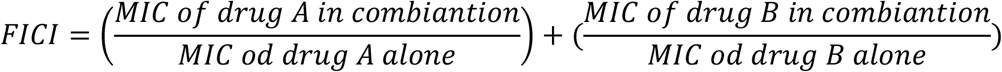

If the value of FICI is ≤ 0.5, the interaction was considered to be synergistic (65). The ZIP score was calculated using synergy finder (24). A ZIP score of > 10 indicates synergy between the two drugs.

### Evolution and Whole Genome Sequencing

Wild-type (WT) cells were evolved under the combined treatment of RIF and CEF. Initially, WT cells were grown up to 0.4 OD_600_. 25 μL of these cells were added in 5 mL of LB containing: No drug; 0.06 μg/mL of RIF with 2.56 μg/mL of CEF; 0.12 μg/mL of RIF with 2.56 μg/mL of CEF; 0.06 μg/mL of RIF with 5.12 μg/mL of CEF. The cultures were allowed to grow overnight. The next day, 25 μL of the overnight cultures were transferred in fresh tubes containing 5 mL of LB with the same conditions. This designated the first passage. All cultures were evolved for 10 passages. Cells from each passage were stored as glycerol stocks. For experiments, the frozen stocks were streaked on LB agar plates and a representative single colony was picked from each passage and analyzed for their RIF and CEF sensitivities. These single colonies were again stored as glycerol stocks. Chromosomal DNA was extracted from the selected single colonies using Qiagen DNA extraction kit and was sent for Whole Genome Sequencing. Sequencing was done using the Illumina platform at the Microbial Genome Sequencing Center (MiGS, Pittsburgh). The results were trimmed, mapped, and aligned with reference WT (NC_000964.3) genome sequence using CLC genomics workbench.

### Disc diffusion assay

Drug susceptibilities of the mutants were screened by determining the zone of inhibition using a disc diffusion assay. Cultures were grown up to ∼0.4 OD_600_. 100 μL of this culture was mixed with 4 mL of top agar (0.75% agar). Top agar was kept at 50°C to prevent it from solidifying. The mix of agar and culture was poured onto a 15 mL LB agar (1.5 %) plates. This was allowed to air-dry for 30 min. A 6 mm Whatmann paper filter disc was then put on the top agar. The required amount of drug was added on the disc immediately. The plates were incubated overnight at 37°C. The diameter of the clear zone of inhibition/low density growth (ZOI/ZOLD) was measured the next day. For all histograms, the Y-axis starts from 6 mm which is the disc diameter. For experiments with GlcNAc supplementation, 20 mM GlcNAc was added in both the top agar and LB agar plates. For strains having the inducible promoter P_*spac(hy)*_, the agar was made with 1 mM IPTG. Amount of drugs used on the disc: CEF – 25 μg; RIF – 25 μg; Oxacillin – 3 μg; Ampicillin – 15 μg; Penicillin G – 20U; Nisin – 100 μg; Vancomycin-10 μg; Fosfomycin-75 μg; Chloramphenicol – 8 μg; Triclosan – 5 μg; Paraquat – 8 μL from 10 mM stock.

### Real-time PCR

Gene expression was determined by real-time PCR using primers mentioned in Supplementary Table 2. Cultures were grown up to ∼0.4 OD_600_. RNA was purified from 1.5 mL of cells using the RNeasy Kit from Qiagen as per the manufacturer’s instructions. The isolated RNA was then given a DNase treatment with TURBO DNA-free Kit (Invitrogen, REF AM1907). Approximately 15 μg of RNA was incubated with 2 μL DNAse and 2 μL Buffer at 37°C for 15 min followed by a 5 min incubation with the DNAse inactivating agent. The samples were then centrifuged at 8000 rpm for 3 min and the supernatant was collected in a fresh micro centrifuge tube. cDNA was prepared with 2 μg of the treated RNA in 20 μL total volume of reaction mix using High-capacity cDNA reverse transcription kit from Applied Biosystems (REF 4368814). The cDNA was further diluted by 1:10 to obtain a final concentration of 10 ng/μL. The gene expression levels were measured using 10 ng of cDNA using 0.5 μM of gene specific primers and 1X SYBR green master mix (Applied Biosystems; REF A25742) in Step-One plus from Applied Biosystems. *gyrA* was used as an internal control. Gene expression values (2^-Δct^) were plotted after normalization with *gyrA*.

### Metabolite Extraction

Metabolomics experiments were done according to previously published work (66, 67). Both wild-type and mutant strains were first grown in 5 ml LB broth (BD Difco™) media at 30 °C for 12 hrs. and then diluted 1:50 in 40 ml media (in triplicates) to grow at 37°C. Mid-log phase cultures with OD_600_ 0.4 were pelleted down and quenched by resuspending in 700 μl of a precooled 40%:40%:20% mixture of acetonitrile, methanol, and water. To extract metabolites, cells were lysed using 0.1 mm Zirconia beads and Precellys homogenizer (Bertin Instruments). Lysates were centrifuged at 12,000 RPM for 8 min at 37°C and cleared by passing through 0.22 μm Spin-X tube filters (Sigma–Aldrich).

### Liquid Chromatography and Mass Spectrometry

2 μl of extracted metabolite samples were separated on a Cogent Diamond Hydride Type C Column of 1200 liquid chromatography (Agilent) which was coupled to an Agilent Accurate-Mass 6220 Time-of-Flight spectrometer. For different classes of metabolites, two types of solvents were used: Solvent A (H_2_O + 0.2% formic acid) and Solvent B (acetonitrile + 0.2% formic acid). Gradient was 0-2 min, 85% B; 3-5 min, 80% B; 6-7 min, 75% B; 8-9 min, 70% B; 10-11.1 min, 50% B; 11.1-14 min 20% B; 14.1-24 min 5% B with a 10 min re-equilibration period at 85% B at a flow rate of 0.4 ml/min. For dynamic mass axis calibration, a reference mass solution was continuously injected from the isocratic pump. Ion abundances of different metabolites were determined using Profinder 8.0. Log_2_ fold changes were calculated with respect to the abundances in the wild-type strain.

### Statistical analysis

All the experiments were performed with a minimum of 3 biological replicates. One-way ANOVA was used to calculate the statistical significance. Tukey’s comparison test was used to determine significance between all the strains. *P*-value cut-offs have been mentioned in the figure legends. Different letters represent data which are significantly different. Same letter represents mean values which are not statistically different. Significance between two strains was determined using student’s t-test.

## Acknowledgments

Research reported in this publication was supported by the National Institutes of Health under award number R35GM122461 to JDH and U19AI162584 and R25140472 to KYR. The content is solely the responsibility of the authors and does not necessarily represent the official views of the National Institutes of Health.

## Graphical Abstract

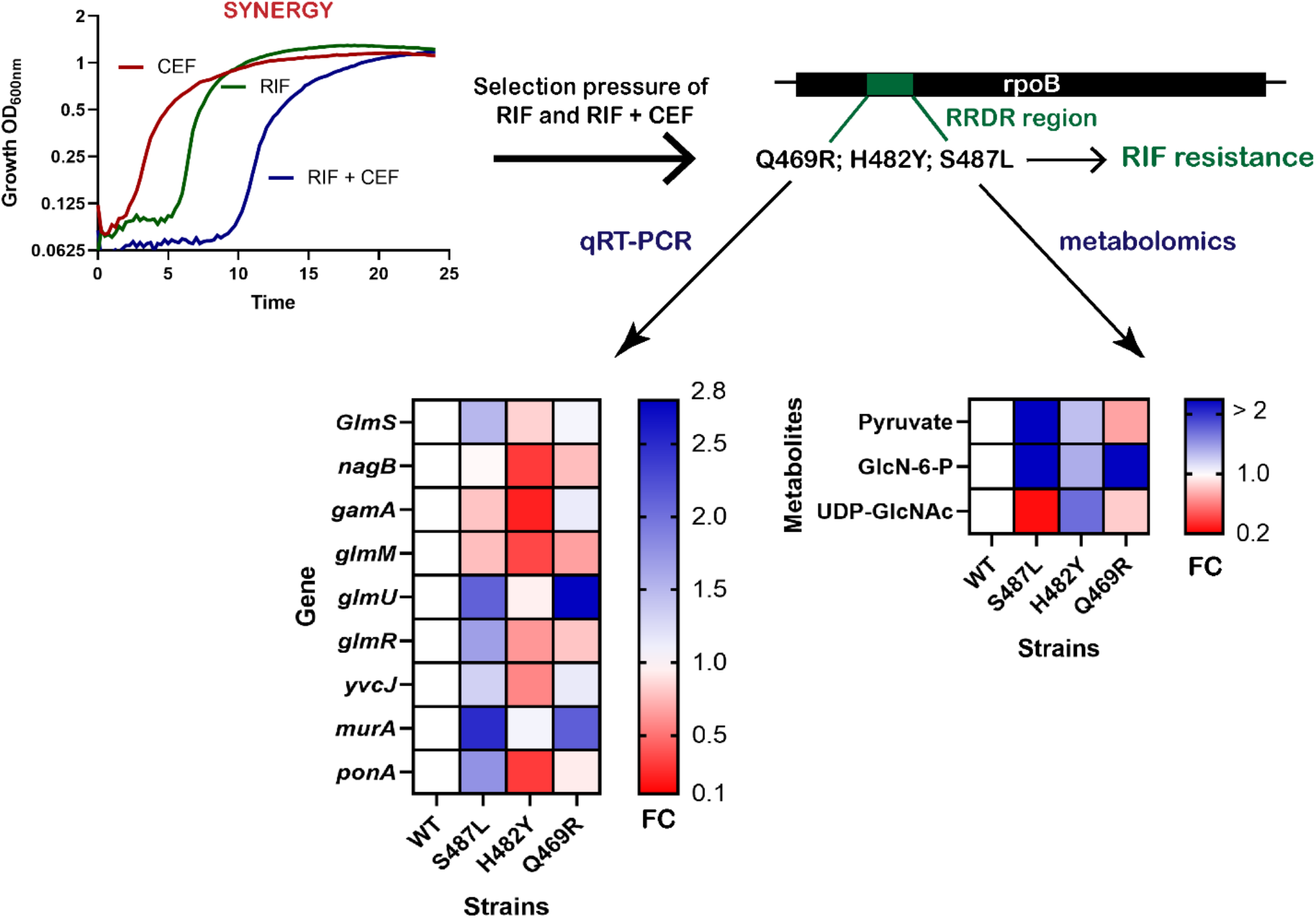

