## Supplementary material for "Mutations in *rpoB* that confer rifampicin resistance can alter levels of peptidoglycan precursors and affect β-lactam susceptibility": SI materials

Short title: *rpoB* mutants and β-lactam susceptibility

Yesha Patel[a], Vijay Soni[b], Kyu Y. Rhee[b], John D. Helmann[a*]

[a]Department of Microbiology, Cornell University, Ithaca NY 14853-8101

[b]Department of Medicine, Division of Infectious Diseases, Weill Cornell Medicine, New York NY 10021-5608

ORCID IDS: orcid.org/0000-0001-9888-9888 (YP), orcid.org/0000-0002-3395-7429 (VS), orcid.org/ 0000-0003-4582-2895 (KYR), orcid.org/0000-0002-3832-3249 (JDH)

Co-author

**Table 1** Bacterial strains used in the study

| Strain number | Strain name | Additional details |
|---|---|---|
| HB26336 | Wild type *B. subtilis* 168 | |
| HB26337 | 0.06R + 2.56C: 4th passage | Evolution; S487L (Strain A) |
| HB26338 | 0.12R + 2.56C: 4th passage | Evolution; S487L (Strain B) |
| HB26378 | 0.06R + 5.12C: 4th passage | Evolution; P520L (Strain C) |
| HB26266 | Q469R | Picked up from agar plates containing RIF |
| HB26341 | H482Y | Picked up from agar plates containing RIF |
| HB28012 | *ΔglmR* | *glmR::null* in HB26336 |
| HB28018 | S487L *ΔglmR* | *glmR::null* in HB26337 |
| HB28016 | H482Y *ΔglmR* | *glmR::null* in HB26341 |
| HB26492 | WT- $P_{spac(hy)}$- glmM | lacA::$P_{spac(hy)}$- glmM in WT using pPL82 |
| HB26498 | S487L- $P_{spac(hy)}$- glmM | lacA::$P_{spac(hy)}$- glmM in HB26337 using pPL82 |
| HB26496 | H482Y- $P_{spac(hy)}$- glmM | lacA::$P_{spac(hy)}$- glmM in HB26341 using pPL82 |
| HB26494 | Q469R- $P_{spac(hy)}$- glmM | lacA::$P_{spac(hy)}$- glmM in HB26266 using pPL82 |
| HB26482 | WT- $P_{spac(hy)}$- pgcA* | lacA::$P_{spac(hy)}$- pgcA(G47S) in WT using pPL82 |
| HB26488 | S487L- $P_{spac(hy)}$- pgcA* | lacA::$P_{spac(hy)}$- pgcA(G47S) in HB26337 using pPL82 |
| HB26486 | H482Y- $P_{spac(hy)}$- pgcA* | lacA::$P_{spac(hy)}$- pgcA(G47S) in HB26341 using pPL82 |
| HB26484 | Q469R- $P_{spac(hy)}$- pgcA* | lacA::$P_{spac(hy)}$- pgcA(G47S) in HB26266 using pPL82 |
| HB26472 | WT- $P_{spac(hy)}$- pgcA | lacA::$P_{spac(hy)}$- pgcA in WT using pPL82 |
| HB26478 | S487L- $P_{spac(hy)}$- pgcA | lacA::$P_{spac(hy)}$- pgcA in HB26337 using pPL82 |
| HB26476 | H482Y- $P_{spac(hy)}$- pgcA | lacA::$P_{spac(hy)}$- pgcA in HB26341 using pPL82 |
| HB26474 | Q469R- $P_{spac(hy)}$- pgcA | lacA::$P_{spac(hy)}$- pgcA in HB26266 using pPL82 |
| HB28124 | *ΔgamA* | *gamA::mls* in HB26336 |

| | | |
|---|---|---|
| HB28127 | S487L *ΔgamA* | *gamA::mls* in HB26337 |
| HB28126 | H482Y *ΔgamA* | *gamA::mls* in HB26341 |
| HB28125 | *Q469R ΔgamA* | *gamA::mls* in HB26266 |

**Table 2** Primers used in the study

| Primer Number | Primer Name | Primer Sequence |
|---|---|---|
| 9284 | rpoB-FP | GATGAAGTTTCCGTCGTTCAA |
| 9285 | rpoB-RP | GAAAATGCGTTCGCAGAATAG |
| 9286 | rpoB-seq-CI | CTTCTCCAGAACCAATTCCGT |
| 9728 | gamA-FP check | CAAGTCCGGCTACACCTTCT |
| 9729 | gamA-RP check | TGCAGAGCCAGTCTCACAGT |
| 9610 | glmS-qFP | AGAAAAAGGACGCATTGCAG |
| 9611 | glmS-qRP | TGAGCGTTCAGATAGCTTGG |
| 9614 | gamA-qFP | TCGGCTTGTATAAGCAGTTGA |
| 9615 | gamA-qRP | ACTTTGCGGATGAGATGGAG |
| 9616 | nagB-qFP | GCCTGCACCAAACTGAGAAT |
| 9617 | nagB-qRP | AAGTGATAGCTGTTCGGGTCA |
| 9612 | glmR-qFP | AATGTGCTTGCCGCTTTATC |
| 9613 | glmR-qRP | TCAAATTCCCGAGAGAATGG |
| 9608 | yvcJ-qFP | CCTTCATTGCTTCCGAAGTT |
| 9609 | yvcJ-qRP | TCAATCAGCCGGTCAAAAA |
| 9606 | glmM-qFP | CAAACAACGTCCAAAAGTGC |
| 9607 | glmM-qRP | ACTTCTGCGCCAATGGATAA |
| 9604 | glmU-qFP | GGTGCGGAAGAAGTGAAAAA |
| 9605 | glmU-qRP | AAGAAATGGCTGTGCCTGTT |
| 9618 | murA-qFP | CACAGACGCACGCATTTTAC |
| 9619 | murA-qRP | GGAGCAGATGTGCATTTTGA |
| 9620 | ponA-qFP | CGGAAAAGCGGCGGAAGAATT |
| 9621 | ponA-qRP | TTATCCGGGTTTTTGACCGGG |
| 8726 | gyrA-qFP | GGCGGCCATGCGTTATACAG |
| 8727 | gyrA-qRP | GCCATACCTACCGCAATGCC |

A

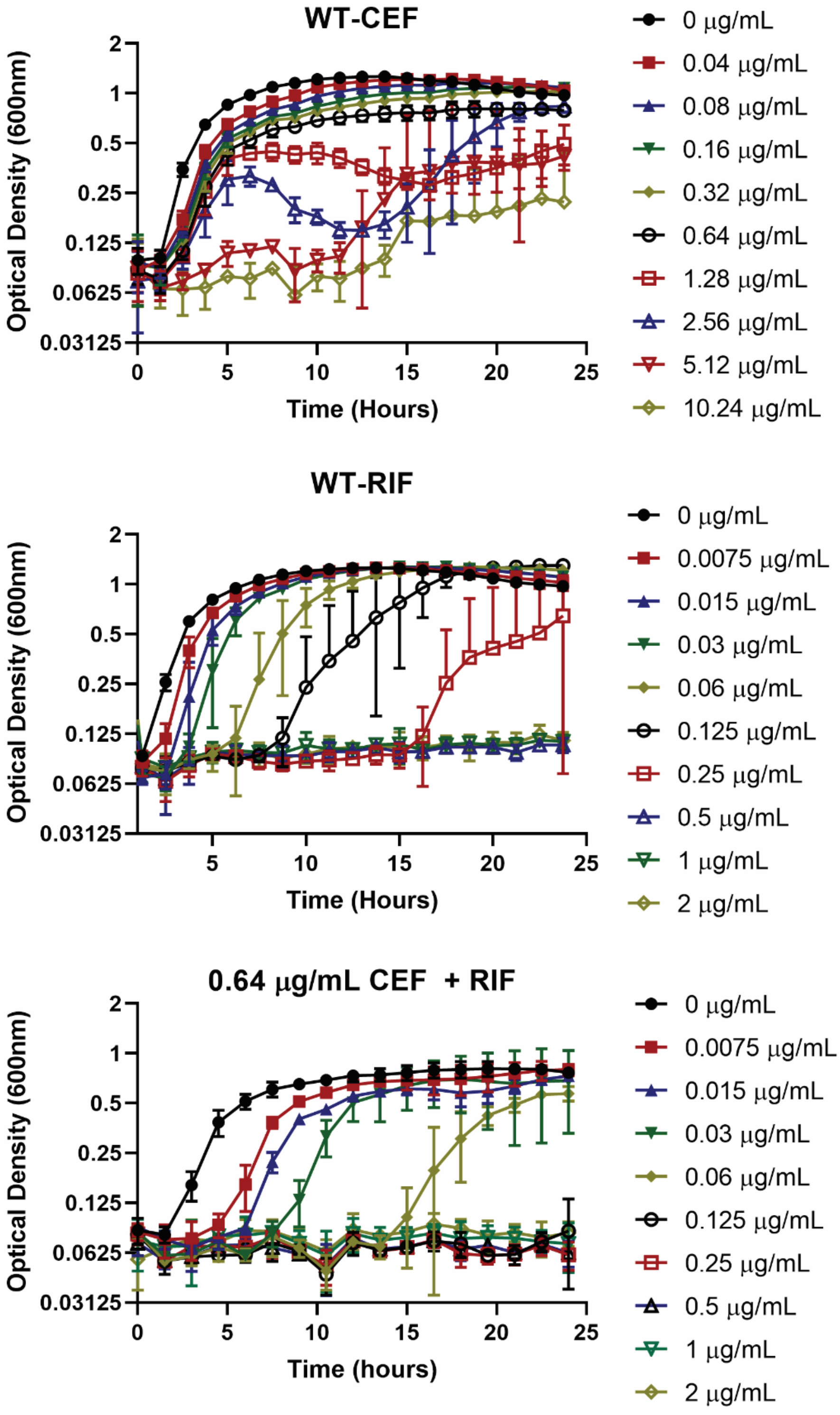
WT-CEF
Optical Density (600nm)
2
1
0.5
0.25
0.125
0.0625
0.03125
0
5
10
15
20
25
Time (Hours)
0 µg/mL
0.04 µg/mL
0.08 µg/mL
0.16 µg/mL
0.32 µg/mL
0.64 µg/mL
1.28 µg/mL
2.56 µg/mL
5.12 µg/mL
10.24 µg/mL
B
WT-RIF
Optical Density (600nm)
2
1
0.5
0.25
0.125
0.0625
0.03125
5
10
15
20
25
Time (Hours)
0 µg/mL
0.0075 µg/mL
0.015 µg/mL
0.03 µg/mL
0.06 µg/mL
0.125 µg/mL
0.25 µg/mL
0.5 µg/mL
1 µg/mL
2 µg/mL
C
0.64 µg/mL CEF + RIF
Optical Density (600nm)
2
1
0.5
0.25
0.125
0.0625
0.03125
0
5
10
15
20
25
Time (hours)
0 µg/mL
0.0075 µg/mL
0.015 µg/mL
0.03 µg/mL
0.06 µg/mL
0.125 µg/mL
0.25 µg/mL
0.5 µg/mL
1 µg/mL
2 µg/mL

**Figure 1: Minimum inhibitory concentrations (MIC) determined by growth kinetics.** Cell density was monitored after treatment with (A) CEF, (B) RIF, or (C) combination of 0.64 µg/mL CEF with RIF at concentrations mentioned in the legend. MIC is defined as >90% growth inhibition compared to untreated control after 8 hrs of treatment.

**Table 3** Synergy determination between RIF and CEF by ZIP score

| CEF (μg/mL) | RIF (μg/mL) | ZIP score |
|---|---|---|
| 0 | 0 | 0.00 |
| 0 | 0.0075 | 0.00 |
| 0 | 0.015 | 0.00 |
| 0 | 0.03 | 0.00 |
| 0 | 0.06 | 0.00 |
| 0 | 0.125 | 0.00 |
| 0 | 0.25 | 0.00 |
| 0 | 0.5 | 0.00 |
| 0 | 1 | 0.00 |
| 0 | 2 | 0.00 |
| 0.04 | 0 | 0.00 |
| 0.04 | 0.0075 | **14.43** |
| 0.04 | 0.015 | **20.23** |
| 0.04 | 0.03 | **25.75** |
| 0.04 | 0.06 | **61.68** |
| 0.04 | 0.125 | **36.48** |
| 0.04 | 0.25 | 6.83 |
| 0.04 | 0.5 | -1.07 |
| 0.04 | 1 | -1.55 |
| 0.04 | 2 | -2.91 |
| 0.08 | 0 | 0.00 |
| 0.08 | 0.0075 | **16.41** |
| 0.08 | 0.015 | **24.05** |
| 0.08 | 0.03 | **34.72** |
| 0.08 | 0.06 | **69.12** |
| 0.08 | 0.125 | **36.07** |
| 0.08 | 0.25 | 7.31 |
| 0.08 | 0.5 | -0.29 |
| 0.08 | 1 | -0.74 |
| 0.08 | 2 | -2.10 |
| 0.16 | 0 | 0.00 |
| 0.16 | 0.0075 | 14.88 |
| 0.16 | 0.015 | 22.77 |
| 0.16 | 0.03 | 36.11 |
| 0.16 | 0.06 | 65.66 |
| 0.16 | 0.125 | 33.41 |
| 0.16 | 0.25 | 6.57 |
| 0.16 | 0.5 | -0.46 |

| | | |
|---|---|---|
| 0.16 | 1 | -0.88 |
| 0.16 | 2 | -2.23 |
| 0.32 | 0 | 0.00 |
| 0.32 | 0.0075 | 8.93 |
| 0.32 | 0.015 | 15.95 |
| 0.32 | 0.03 | 33.93 |
| 0.32 | 0.06 | 55.81 |
| 0.32 | 0.125 | 27.66 |
| 0.32 | 0.25 | 4.42 |
| 0.32 | 0.5 | -1.54 |
| 0.32 | 1 | -2.06 |
| 0.32 | 2 | -3.39 |
| 0.64 | 0 | 0.00 |
| 0.64 | 0.0075 | 3.89 |
| 0.64 | 0.015 | 10.36 |
| 0.64 | 0.03 | 35.58 |
| 0.64 | 0.06 | 43.14 |
| 0.64 | 0.125 | 20.19 |
| 0.64 | 0.25 | 2.24 |
| 0.64 | 0.5 | -1.83 |
| 0.64 | 1 | -2.91 |
| 0.64 | 2 | -3.00 |
| 1.28 | 0 | 0.00 |
| 1.28 | 0.0075 | 2.31 |
| 1.28 | 0.015 | 3.88 |
| 1.28 | 0.03 | 24.21 |
| 1.28 | 0.06 | 27.61 |
| 1.28 | 0.125 | 12.03 |
| 1.28 | 0.25 | -0.12 |
| 1.28 | 0.5 | -2.70 |
| 1.28 | 1 | -3.57 |
| 1.28 | 2 | -3.82 |
| 2.56 | 0 | 0.00 |
| 2.56 | 0.0075 | 4.44 |
| 2.56 | 0.015 | 4.89 |
| 2.56 | 0.03 | 11.85 |
| 2.56 | 0.06 | 14.26 |
| 2.56 | 0.125 | 4.99 |
| 2.56 | 0.25 | -2.26 |
| 2.56 | 0.5 | -3.81 |
| 2.56 | 1 | -4.26 |
| 2.56 | 2 | -4.75 |
| 5.12 | 0 | 0.00 |

| | | |
|---|---|---|
| 5.12 | 0.0075 | 4.36 |
| 5.12 | 0.015 | 5.03 |
| 5.12 | 0.03 | 4.64 |
| 5.12 | 0.06 | 5.20 |
| 5.12 | 0.125 | 0.19 |
| 5.12 | 0.25 | -3.71 |
| 5.12 | 0.5 | -4.65 |
| 5.12 | 1 | -5.07 |
| 5.12 | 2 | -5.69 |
| 10.24 | 0 | 0.00 |
| 10.24 | 0.0075 | -1.54 |
| 10.24 | 0.015 | -0.77 |
| 10.24 | 0.03 | -1.22 |
| 10.24 | 0.06 | -0.01 |
| 10.24 | 0.125 | -2.54 |
| 10.24 | 0.25 | -4.36 |
| 10.24 | 0.5 | -5.00 |
| 10.24 | 1 | -5.37 |
| 10.24 | 2 | -5.99 |

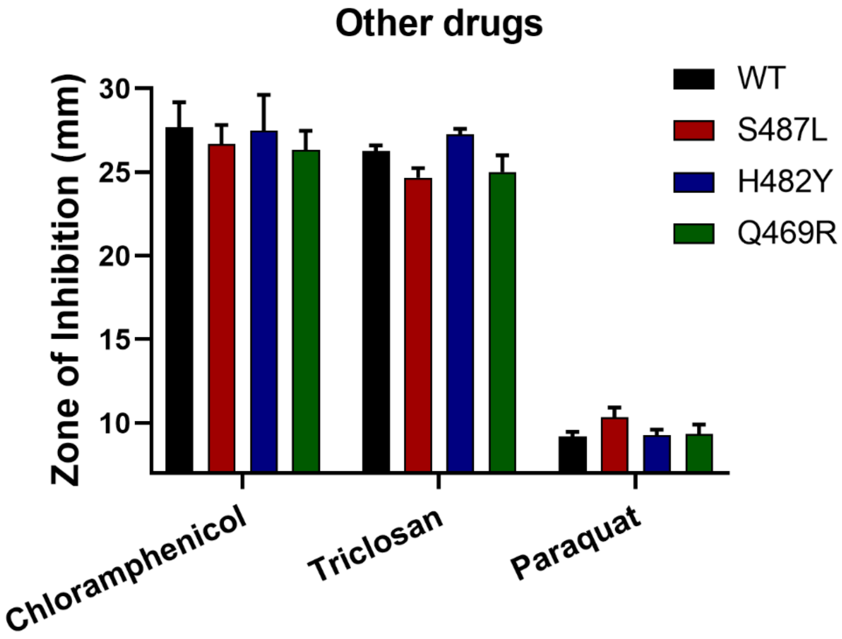


**Figure 2 Drug susceptibilities of *rpoB* mutants as measured by zone of inhibition against drugs that do not directly target cell wall synthesis**. Chloramphenicol inhibits protein synthesis; triclosan inhibits fatty acid synthesis and paraquat generates ROS toxicity in the cells.

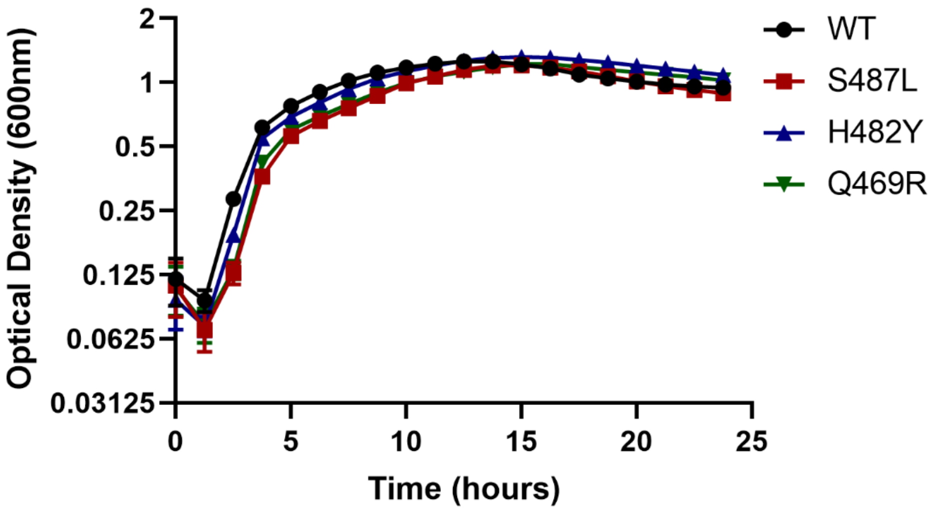


**Figure 3 Growth kinetics of *rpoB* mutants in LB medium in the absence of any drug treatment.**

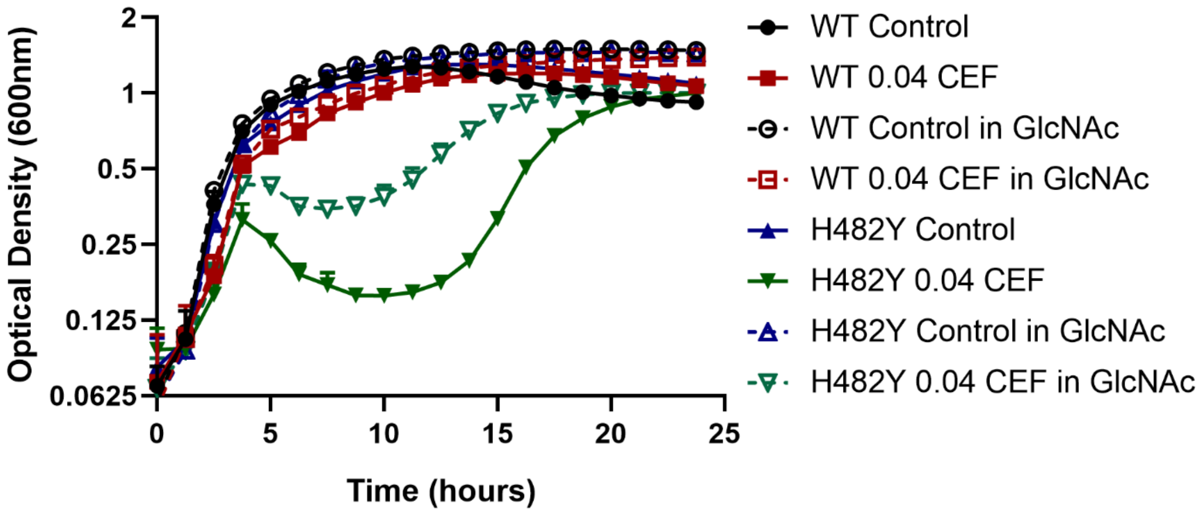


**Figure 4 Growth kinetics of WT and H482Y mutant.** Growth was monitored in LB (Control), LB with 0.04 μg/mL of CEF, LB supplemented with GlcNAc, LB supplemented with GlcNAc and 0.04 μg/mL of CEF

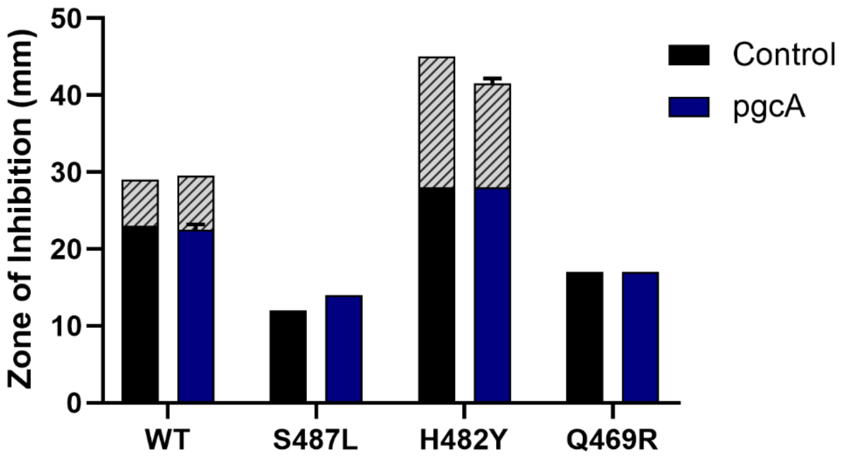


**Supplementary Figure 5 CEF susceptibility as measured by zone of inhibition assay for WT and *rpoB* mutants.** Black Control bars represent the CEF sensitivity of the mutants. Blue *pgcA* bars represents the CEF sensitivity of the respective strains with the *pgcA* gene deleted.
